# Optokinetic nystagmus reflects perceptual directions in the onset binocular rivalry in Parkinson’s disease

**DOI:** 10.1101/074898

**Authors:** Mana Fujiwara, Catherine Ding, Lisandro Kaunitz, Julie C Stout, Dominic Thyagarajan, Naotsugu Tsuchiya

**Affiliations:** School of Psychological Sciences, Faculty of Medicine, Nursing and Health Sciences, Monash University, Melbourne, Victoria, Australia; Araya Brain Imaging, Tokyo, Japan; Department of Neurosciences, Southern Clinical School, Monash Health, Melbourne, Victoria, Australia; Research Center for Advanced Science and Technology (RCAST), The University of Tokyo, Tokyo, Japan; Monash Institute of Cognitive and Clinical Neurosciences, Monash University, Melbourne, Victoria, Australia

**Author notes:** Corresponding authors: (NT) and (DT). These authors contributed equally to this work (Mana Fujiwara, Catherine Ding).

## Abstract

Optokinetic nystagmus (OKN), the reflexive eye movements evoked by a moving field, has recently gained interest among researchers as a useful tool to assess conscious perception. When conscious perception and stimulus are dissociated, such as in binocular rivalry —when dissimilar images are simultaneously presented to each eye and perception alternates between the two images over time — OKN correlates with perception rather than with the physical direction of the moving field. While this relationship is well established in healthy subjects it is yet unclear whether it also generalizes to clinical populations, for example, patients with Parkinson’s disease. Parkinson’s disease is a motor disorder, causing tremor, slow movements and rigidity. It may also be associated with oculomotor deficits, such as impaired saccades and smooth pursuit eye movements. Here, we employed short-duration, onset binocular rivalry (2 s trial of stimulus presentation followed by 1 s inter-trial interval) with moving grating stimuli to assess OKN in Parkinson’s disease patients (N=39) and controls (N=29) of a similar age. Each trial was either non-rivalrous (same stimuli presented to both eyes) or rivalrous, as in binocular rivalry. We analyzed OKN to discriminate direction of stimulus and perception on a trial-by-trial basis. OKN reflected conscious perceptions in both groups. Treatment with anti-Parkinson drugs and deep brain stimulation improved motor ability of patients assessed by a standard scale of Parkinson’s disease, but did not impact on OKN. Furthermore, OKN-based measures were robust and their latencies were shorter than manual button-based measures in all subjects, regardless of stimulus condition. Our findings suggest that OKN can be used as an indicator of conscious perception in binocular rivalry even in Parkinson’s disease patients in whom impaired manual dexterity may render button-press reports less reliable.

## Introduction

By the time motor deficits are apparent in Parkinson’s disease patients (PD), more than 70% of dopamine-generating neurons are lost (1). As the neurodegeneration progresses, so does motor disability, manifesting in slowed and small movements (bradykinesia/hypokinesia) and muscle rigidity. Neurodegeneration also affects saccades (2–8), smooth pursuit eye movements (2–6,8,9) and visual perception (10). Reduced mobility creates difficulties for perceptual research on PD patients because impaired motor dexterity renders manual responses unreliable.

Recently, optokinetic nystagmus (OKN), a reflexive eye-movement evoked by a moving field, has been gaining traction among researchers as a tool to assess perception, because it can reliably signal conscious perception under certain experimental paradigms (11,12). In particular, under binocular rivalry, where dissimilar images are presented to each eye and subjects continuously report changes in their perception from one image to the other, a tight correlation between OKN and perceptual switches has been established primarily in healthy adult humans and macaque monkeys (11,13–18) (19). However, it remains largely unknown whether such a correlation exists across all populations, including those with brain diseases. The question is of considerable scientific and clinical importance if OKN is to be used as convenient ‘readout’ of perception in clinical populations, such as people who have impaired motor responses due to PD.

To use the OKN as a “readout” of perception in PD, the OKN of patients must be intact, and must reflect perception. However, the effect of PD on the OKN is controversial, some studies reporting impaired horizontal OKN in PD patients (6,7,9) while others claim it is intact (3–5,8,20,21). Moreover, there are no studies on the OKN and perception in PD patients during binocular rivalry. Here, we used the binocular rivalry paradigm to investigate OKN as a “readout” of perception in PD patients. We compared OKN and button press reports in a short-duration ‘onset rivalry’ paradigm (22). In this paradigm, rivalrous trials (in which stimuli travelling in opposite directions are projected to each eye) are intermixed with non-rivalrous trials (in which the same stimuli is projected to both eyes). Non-rivalrous trials provide a baseline estimate of the speed and accuracy of OKN and button press. We hypothesized that manual and ocular responses would be slowed down in PD patients compared to controls as well as within patients when they are off-treatment than on-treatment. Our study design allows us to qualitatively compare the accuracy and latency of manual and ocular responses in rivalrous condition compared against control non-rivalrous condition. We assessed the effects of disease with between-subjects comparisons (PD patients vs similar age control) as well as the effects of treatment of PD with within-subject comparisons (on vs off treatment).

## Materials and Methods

### Subjects

Thirty-nine PD patients and 29 age-similar normal controls participated in this study. We tested a further 9 PD patients, but did not include them because they did not complete two testing sessions. Seventeen patients had been implanted with electrodes for deep brain stimulation (DBS) treatment and the other 22 patients were treated only with medication. We report three kinds of effects: 1) PD — between-subject comparisons of PD patients and controls, 2) medication — within-subject comparisons of patients in on-medication and off-medication states, and 3) DBS — within-subject comparisons of patients in on-DBS and off-DBS states. Thus, all PD patients were tested twice — with and without medication or with and without DBS, in counterbalanced order. In each session, the severity of each patient’s motor deficits was measured using the Movement Disorder Society Unified Parkinson's Disease Rating Scale (MDS-UPDRS) Part III (motor examination). For further details of subject demography see Results section. All subjects gave written informed consent prior to their participation in the study. The study conformed to the Declaration of Helsinki and was approved by the Monash Health Research Ethics Committee (MUHREC 12350B).

### Apparatus

Stimuli were created using Matlab Psychtoolbox (23,24) with Matlab 2013 on a MacBook Pro and displayed on a 23” Tobii TX-300 screen (resolution: 1920 x 1080 pixels, refresh rate 60 Hz). Eye movements were recorded using a Tobii TX-300 eye tracker (Tobii Technology, Danderyd, Sweden) at a sampling rate of 300Hz. Tobii was controlled by Matlab with the software package T2T (http://psy.cns.sissa.it/t2t/About_T2T.html).

Subjects sat in a brightly-lit room on a height-adjustable chair, with their heads stabilized by a chin rest at a distance of 74 cm from the monitor. Each grating stimulus was projected to each eye, through a custom-built stereoscope consisting of four mirrors. Two of the mirrors were transparent to infrared light, allowing the eye-tracker to track eye position whilst restricting each eye to viewing only one grating (18). Subjects responded with both hands using numeric keypads. The button press data was recorded in Matlab at the sampling rate of 60Hz.

### Stimuli

We tested subjects’ OKN with moving sinusoidal gratings. The gratings were presented at the center of each mirror, confined within a square area of 5.34 degrees of visual angle. The gratings had a spatial frequency of 0.27 cycles per degrees of visual angle and a temporal frequency of 6.02 cycles per second. As a result, the gratings drifted at a speed of 22.3 degrees of visual angle per second. Each grating was framed by a square box of a random texture pattern, which facilitated binocular fusion in subjects.

### Experimental procedures

Subjects viewed stimuli through a mirror stereoscope (Fig 1A). Before starting the experiments, we checked that they had achieved binocular fusion. Subjects were instructed to report the dominant direction of motion of the gratings as leftwards with their left hand and as rightwards with their right hand, using the respective keypads (Fig 1A). They were asked to press and hold-down the key as soon as they saw the grating moving left or right. During the 1-s inter-trial interval, they were asked to release the button.

**Fig 1.**
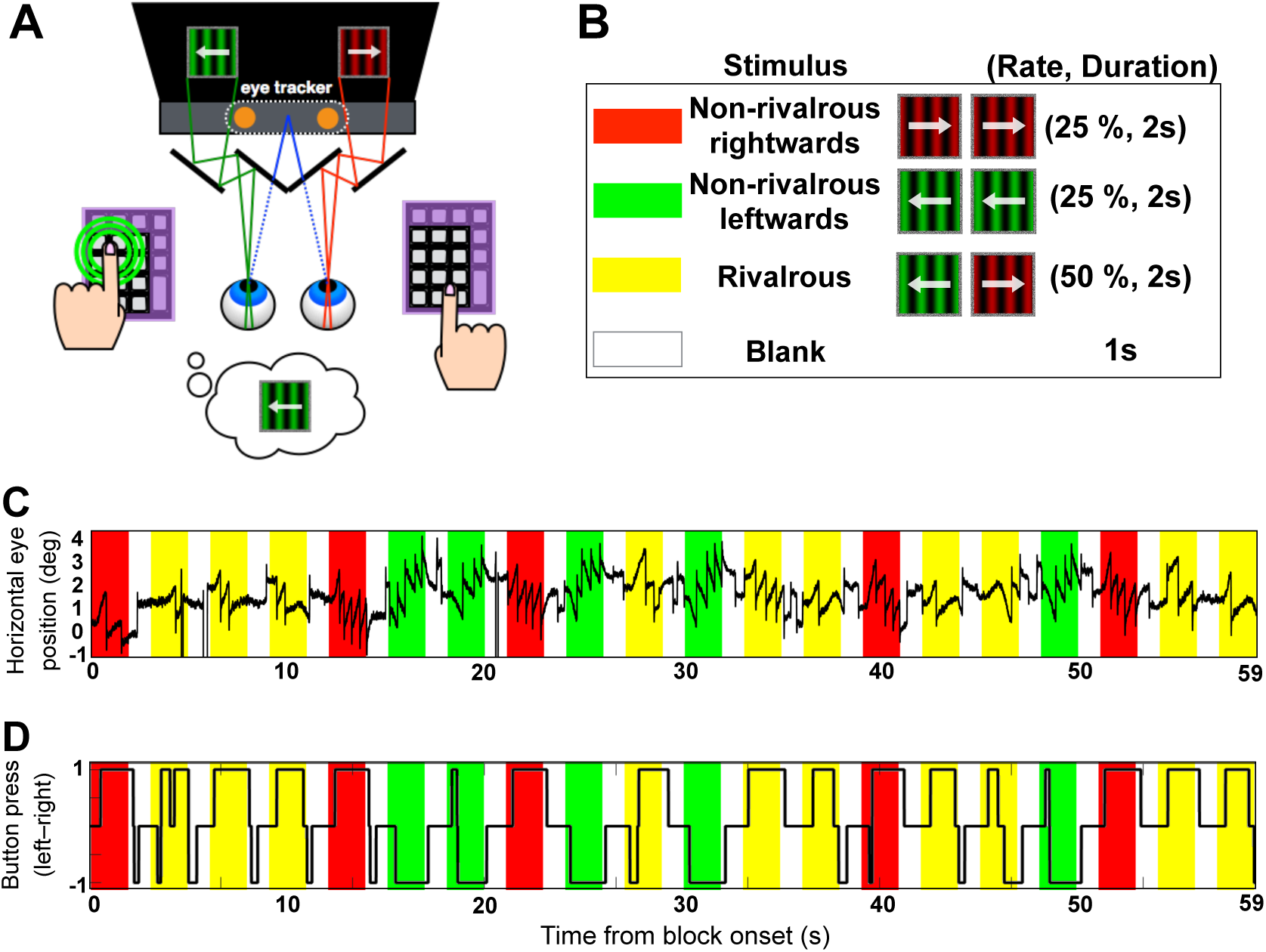
Experimental paradigm. (A) A mirror stereoscope was used to present distinct images to each eye. The mirrors in front of the subject’s eyes were transparent to infrared light, allowing for eye tracking. Subjects were instructed to press a key according to their dominant perception: on the left keypad when the green gratings appeared to go leftwards, and on the right keypad when the red gratings appeared to go rightwards. (B) Each block contained 20 repetitions of a 2-s trial with a 1-s blank interval between trials. Non-rivalrous leftwards, non-rivalrous rightwards or rivalrous trials were randomly intermixed, composing 25 %, 25 % and 50 % of trials in each block, depicted as red, green and yellow in panel B, C, and D, respectively. (C) An exemplar time course of horizontal eye position in one PD patient for 1 block. Center of the stimulus corresponds to the eye position of 0 deg. In non-rivalrous trials (red and green) there is clear OKN with slow phase moving in the same direction as the stimulus, and fast phase (saccades) moving the eyes back. In this block, horizontal eye position was positively biased towards right. Slight bias is expected as we did not provide a fixation point, which reduces OKN. (D) the button press time course from the same block as above. Correspondence between button press report and the OKN can be observed during all trial types, including rivalrous condition in yellow.

Before the main experiment, subjects practiced the task for 4 blocks of 30 s. During practice we confirmed: 1) that the subject pressed just one button, either left or right, and not both at the same time, 2) that button presses reflected the direction of non-rivalrous stimulus in the corresponding non-rivalrous trials, and, 3) that the subject released the button during the blank periods. After each practice run, we plotted the time course of the stimuli and button press on the display (similar to Fig 1D) and proceeded when we were satisfied that the subject understood and followed the instructions.

Before each block of the main experiment, we checked binocular fusion and calibrated the eye-tracker using our custom-written 5-point calibration program that accommodates the mirror setup. After each block, we checked the quality of eye fixation point data and re-calibrated the eye-tracker if less than 80 % of data was validly recorded according to Tobii’s criteria. Eye movements were continuously recorded from the beginning to the end of each block. Each block lasted 60 s and contained 10 non-rivalrous and 10 rivalrous trials, randomly intermixed (Fig 1B, C and D). An experimental session consisted of 8 blocks and lasted approximately 40 minutes (due to a technical error, we failed to record the timing signal in the last block of each session, thus we used first 7 blocks of each session for the analysis). Subjects were allowed to take rests anytime in between blocks if they felt tired or drowsy. When time allowed, subjects were tested with further blocks if recording of eye gaze data was poor in the preceding blocks.

### Behavioral analysis

Before proceeding with eye tracking analyses, we excluded trials and subjects based on the button press data. First, we excluded trials if the button press from the previous trial was not released during the subsequent inter-trial interval, which makes it impossible to define the latency of the button press for the next trial. We also excluded trials where there was either no button press or double button presses (both left and right at the same time) for more than 1 s during the first 2 s of the trial. Second, we rejected subjects if more than half of their rivalrous or non-rivalrous trials were rejected based on the above button press criteria. Table 1 summarizes rejection of trials and subjects.

**Table 1.** Rejection of trials and subjects. Trials were rejected due to the button press (BP) criteria (See Method). Subjects were rejected if more than half of total trials were rejected. Note that controls were tested in one session while PD were tested two sessions, explaining a larger number, but comparable ratio, of rejected trials. “Effect of both” represents effect of both button press and eye movement. “Imbal.” represents imbalance of perceptual direction, meaning that less than 3 trials were classifiable as either dominantly-left or dominantly-right responses.

| Subject Group |  | Control | PD |
| --- | --- | --- | --- |
| Tested subjects |  | 29 | 39 |
| Button Press | Rejected trials due to BP | 1363(29.1%) | 4290(33.1%) |
|  | Rejected subjects | 4(13.8%) | 9(23.1%) |
| Remaining subjects after rejection due to BP |  | 25 | 30 |
| Eye Movement | Rejected trials due to eye movement | 1139(24.3%) | 3538(23.9%) |
|  | Rejected subjects | 0 | 2(7.7%) |
| Effect of both | Rejected trials in total | 1658(35.4%) | 5204(40.1%) |
|  | Rejected subjects | 1(4.0%) | 0 |
| Remaining subjects after rejection due to eye movement |  | 24 | 28 |
| Imbal. | Rejected subjects | 0 | 6(15.4%) |
| Total | Final rejected subjects | 5(17.2%) | 17(43.6%) |
|  | Final subjects | 24 | 22 |

In non-rivalrous condition, accurate button press is objectively defined. To calculate button response accuracy, the direction of the stimulus in each trial (StimDir) was assigned −1 for left and +1 for right. The state of button press at time = t (RawBP(t)), was assigned −1 for left, +1 for right, and 0 for double button press or no button press. Then, we smoothed this RawBP time course with a 100 ms boxcar kernel to obtain the smoothed button press at time t: BP(t). Next, we defined correctness of button press at time = t as C(t), which is 1 if BP(t) * StimDir >= 0.5 and 0 otherwise.

In rivalrous condition, the accuracy of button presses cannot be objectively defined because there is no “correct” answer at any given time. Instead, we labeled each trial either as a dominantly-left or a dominantly-right trial, based on dominant direction of button press over time in that trial. Then we examined the relationship between the dominant percept of a given trial and the button press at a given time of the trial. For this purpose, first we calculated the dominant direction of button press for a given trial (pBP) by averaging BP(t) from t = 0 s to t = 1.5 s. Then, we labeled the trial as dominantly-left (LabelDir = −1) if pBP <0 and dominantly-right (LabelDir = 1) if pBP >0. After labelling each rivalrous trial, we defined button press consistency C(t) at time = t, which is 1 if BP(t) * LabelDir >= 0.5 and 0 otherwise. (When we apply this consistency measure for the non-rivalrous trials, consistency of button press ends up almost identical to button press correctness because >92% of trials were correct and consistent. Thus, we intentionally use the same abbreviation C(t) for consistency/correctness in non-rivalrous trials and consistency in rivalrous trials can be compared.) Ultimately, button response accuracy for a given trial was defined as as the mean C(t), where t ranges from the first button press time at each trial to 1.5 s from the stimulus onset.

Given that there is no objectively “correct” button press in rivalrous condition, we established the first ‘consistent’ button press to allow comparison of button press responses in rivalrous and non-rivalrous conditions. In both conditions, the first consistent button press is defined as the first button press in the direction that the trial is labelled, which occurs when BP(t) * LabelDir >= 0.5. In non-rivalrous trials, first consistent button press is almost the same as the first correct button press. In rivalrous trials, first button press and first consistent button press are the same in trials where the subject did not change their button press, but they can diverge if subjects switch their button press during the trial. For example, consider a trial, in which a subject pressed a left button from 0.5 s to 1 s and switched to right button from 1 s to the end of the trial. In this case, the dominant direction of reported perception (and thus the labelled direction of this trial), first button press time, and first consistent button press time would be “right”, 0.5 s and 1 s, respectively. We calculated the latencies of first button press and of first consistent button press in each trial for both non-rivalrous and rivalrous conditions.

### Eye movement data analysis

We focused our analyses on the velocity of slow-phase OKN, which has been shown to provide a continuous and stable estimate of conscious perception (13–18). To extract the velocity of slow-phase OKN, we performed the following preprocessing to the raw fixation position data.

First, we removed blinks and saccades. We identified blinks as periods of missing fixation position data. We identified saccades with a velocity-based algorithm (25), using a threshold of 6 deg/s in velocity and 1 deg/s^2^ in acceleration. Denoting the i-th removed time period as [R_start_(i), R_end_(i)] and x-coordinate of the fixation position at time=t as F(t), we interpolated F(t) over the time period t = [R_start_(i) − 10 ms, R_end_(i) + 10 ms] with a constant value of F(R_start_(i) − 10 ms). After interpolation, the subsequent fixation position (i.e., F(t) where t >= R_end_(i) + 10 ms) was shifted so that it started from the interpolated position, that is, F(t) was replaced with F(t) − F(R_end_(i) + 10 ms) + F(R_start_(i) − 10 ms) for t >= R_end_(i) + 10 ms. We repeated this procedure until we removed all blinks and saccades in a given block of 60 s. We call the resulting concatenated fixation data as ‘integrated OKN’.

Second, we smoothed the integrated OKN with the same boxcar kernel of 100 ms that we applied to button press time course. We then computed instantaneous velocity of integrated OKN as the difference between neighboring two time points (3.3 ms difference). To obtain a velocity of the slow-phase OKN, we further smoothed the instantaneous velocity with the 100 ms boxcar kernel. Finally, we segmented the time course of the velocity of slow-phase OKN into 3 s epoch, starting from 1 s before the onset of stimuli and ending 2 s after the offset of stimuli. We did not include the first trial of each block in the analysis as we did not record fixation position before the first trial.

We rejected trials if the total length of blinks, saccades and undetected time points by the eye tracker within 2 s of stimulus onset exceeded 1 s. If we rejected more than half of trials for a given session due to either button press criteria (mentioned above) or eye tracking criteria, we excluded the subject from further analysis. After trial rejection by button press and eye movement, if the subject had less than 3 trials classifiable as either dominantly-left or dominantly-right responses, they were also excluded from further analysis. Table 1 shows the breakdown of rejected trials and excluded subjects according to eye tracking criteria.

### Ideal observer analysis on OKN and button press

To compare the relative discriminative qualities of OKN and button press in perceptual report, we employed an ideal observer analysis. That is, we assumed an ideal observer who has access to the distribution of OKN and button press at any given moment.

For this analysis, we first obtained an equal number of left (N_L_) and right (N_R_) trials, by subsampling the trials for side with larger samples. For example, if N_L_ > N_R_, we randomly subsampled N_R_ trials out of N_L_trials. At each time point, we computed the mean velocity of slow-phase OKN from 70 % of the sampled trials. We then defined the midpoint of the two means as the threshold classifier. Using this threshold, we classified the remaining 30 % of sampled trials — trials with a velocity less than the threshold were classified as left trials and those with a velocity greater than threshold were classified right. If the result from this classification was the same as the trial label, we count it as correct. We defined the cross-validation accuracy of the classifier as the mean accuracy for the 30% sampled trials. We repeated this procedure (i.e., from subsampling to cross validation) 10 times and computed the mean cross-validation accuracy at each time point as the discriminability of the OKN.

## Results

In this study, we analyzed the effects of PD and its treatment on behavioural responses and OKN in three separate analyses. For each analyses, we analyzed a specific subset of the data. In the first set of analyses (Fig 2-4), we examined the effects of PD on button press and eye movements in PD patients when they are on treatment (either on-medication or on-DBS), comparing with controls. In the second (Fig 5) and third (Fig 6) set of analyses, we consider the within-subject effects of the medication and DBS, respectively.

**Fig 2.**
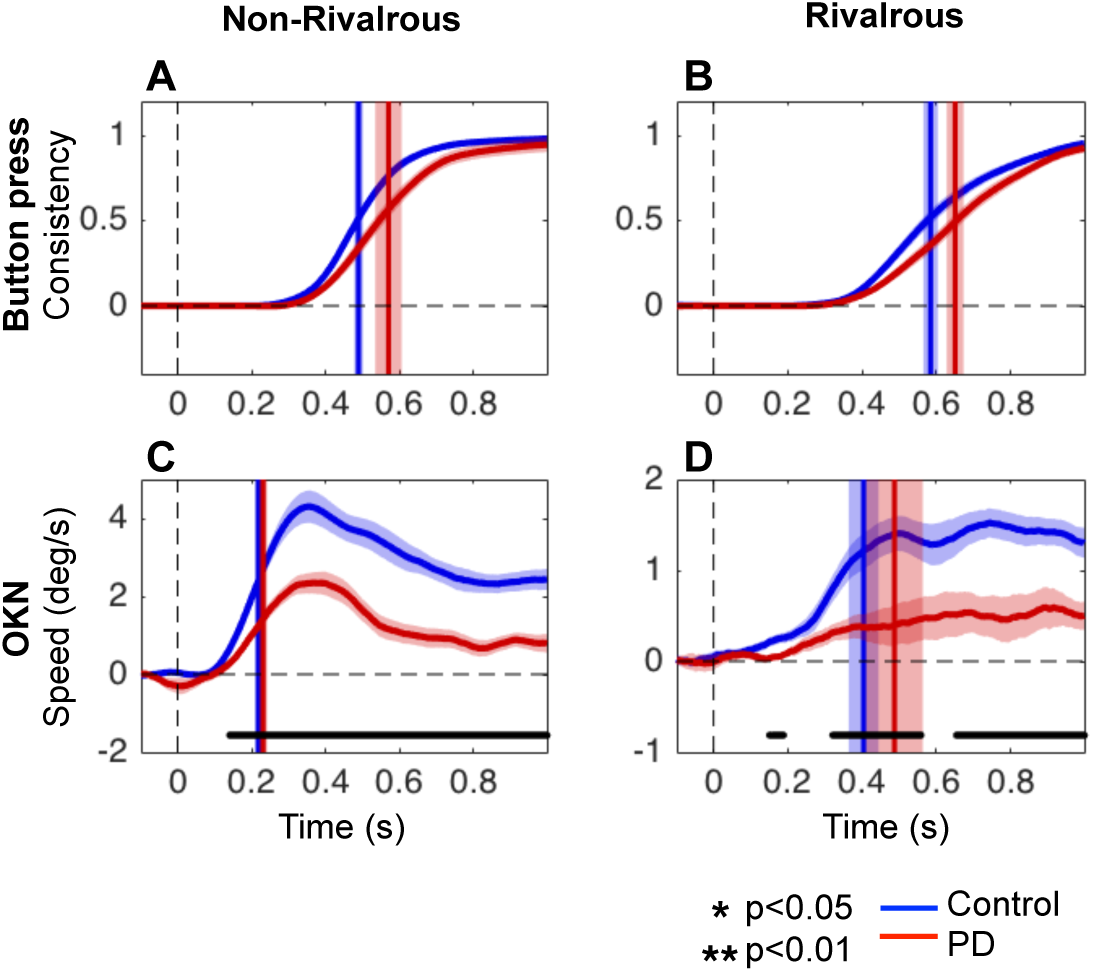
Time course of button press consistency and OKN. The mean time course of button press consistency (A and B) and velocity of slow-phase OKN (C and D) in non-rivalrous (A and C) and rivalrous (B and D) conditions. The vertical lines denote the time of midpoint of the maximum button press consistency or OKN. Black lines at the bottom (only present for C and D) represent the time period where there is a significant difference between PD and controls (unpaired two-tailed t-test, False Discovery Rate adjusted at q=0.05, p<0.05). The data for PD and controls are shown in red and blue, respectively, and the shaded area (when visible) represents SEM across subjects. Note that y-axis for C and D are on different scales.

**Fig 3.**
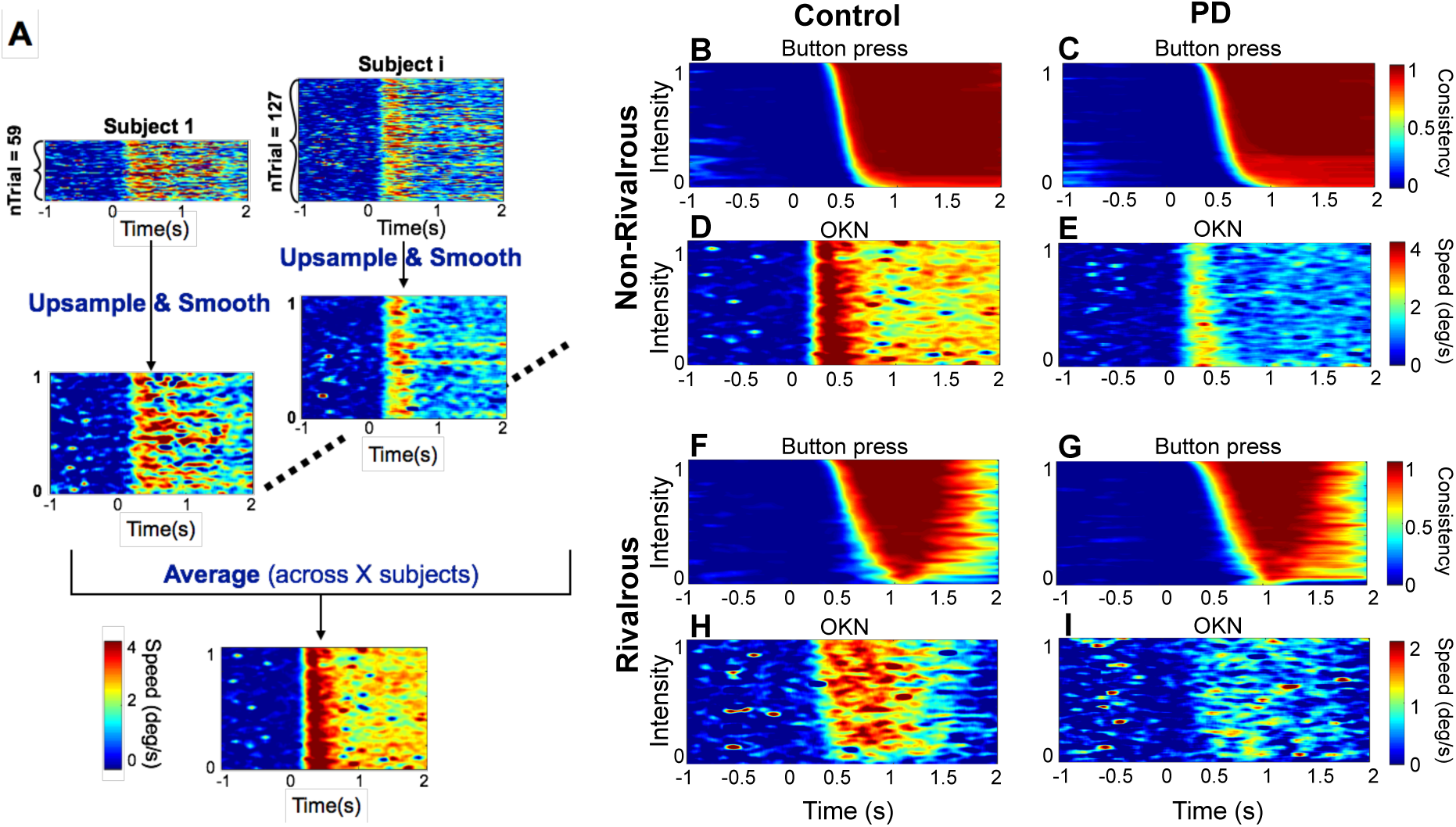
Trial-by-trial image analyses for button press and OKN. A) Preprocessing for image analyses. We first sorted the order of trials according to the button-press consistency, C(t) (see Methods), over each 2 s trial. Next, we stretched the trial x time image along the trial dimension so that we can average the image across subjects who have different numbers of valid trials. Then, we upsampled the image by 1000 points along the trial dimension. Next, we smoothed the image by a boxcar kernel of 31 points along the trial dimension and averaged across subjects. This process was performed for both button press consistency and OKN. B-I) The trial x time images for button-press consistency (B, C, F and G) and OKN (D, E, H and I), across controls (B, D, F, H) and PD (C, E, G, I). Note that button press and OKN velocity was flipped for the left trials (as was done for Fig 2). Button press consistency and OKN speed are denoted according to the color scale key on the right.

**Fig 4.**
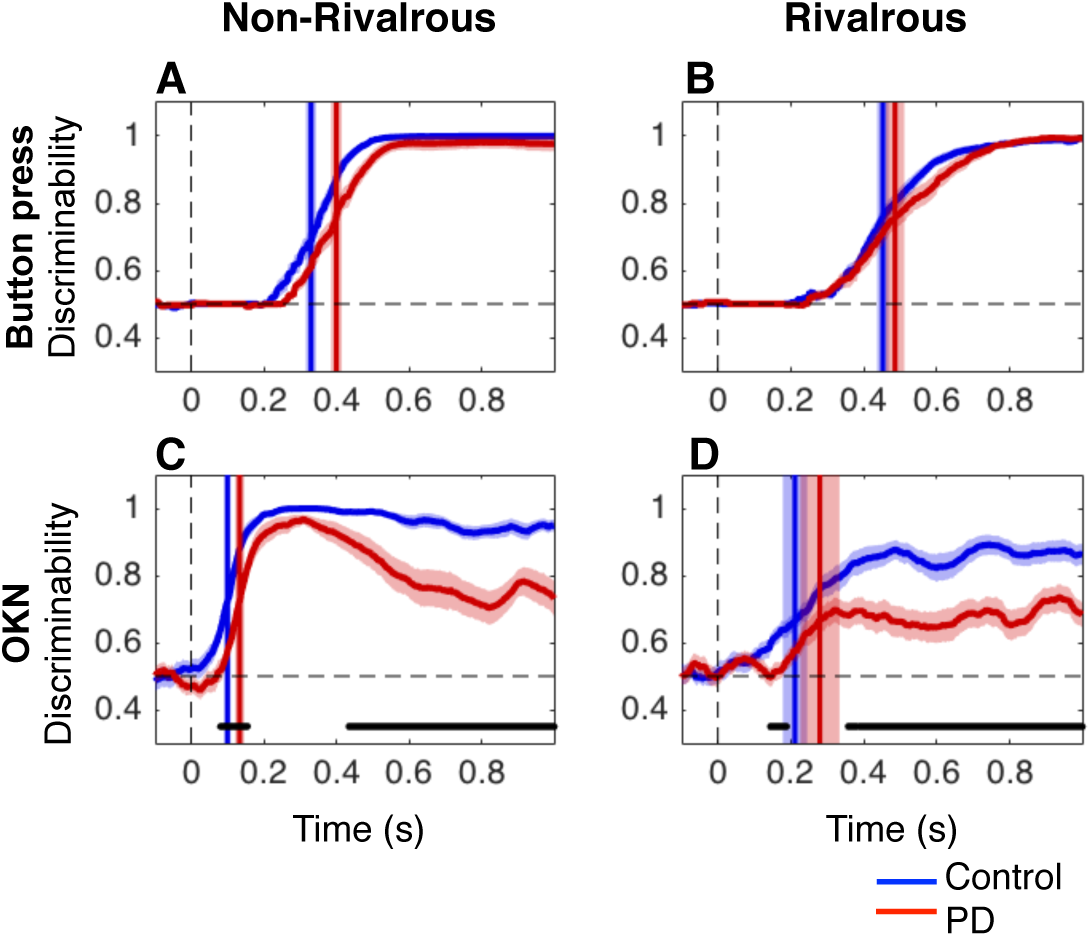
Discriminability analyses. The mean time course of the discriminability of the direction of physical stimuli in non-rivalrous condition (A and C) or dominant percept in rivalrous condition (B and D), measured with button press consistency (A and B) and OKN (C and D). Red and blue lines for PD and controls, respectively. The vertical lines denote time to reach half of the maximum discriminability. Shaded area (when visible) represents SEM across subjects. Black lines at the bottom are significant time points (p<0.05 with FDR q=0.05).

**Fig 5.**
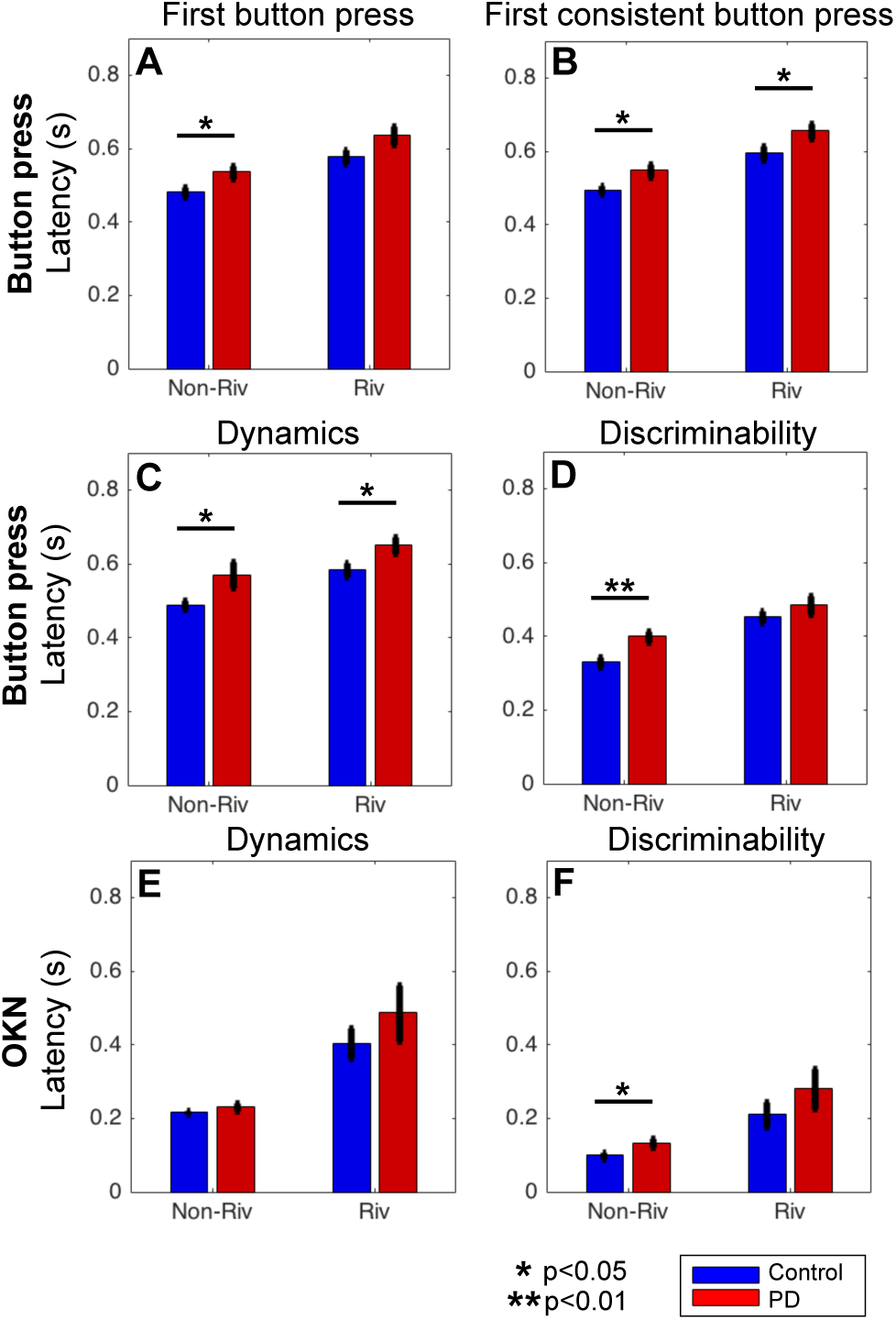
Summary of measures comparing PD (on medication or DBS state N=22) and control (N=24) in non-rivalrous and rivalrous condition. Summary of the results for button press (A-D) and OKN (E and F) for patients with Parkinson’s disease (red) and controls (blue). In each panel, the non-rivalrous (non-riv) condition is on the left and the rivalrous (riv) condition is on the right. Error bars represent SEM across subjects. * and ** indicate significant difference between PD and controls at p<0.05 and p<0.01 respectively (unpaired two-tailed t-test).

**Fig 6.**
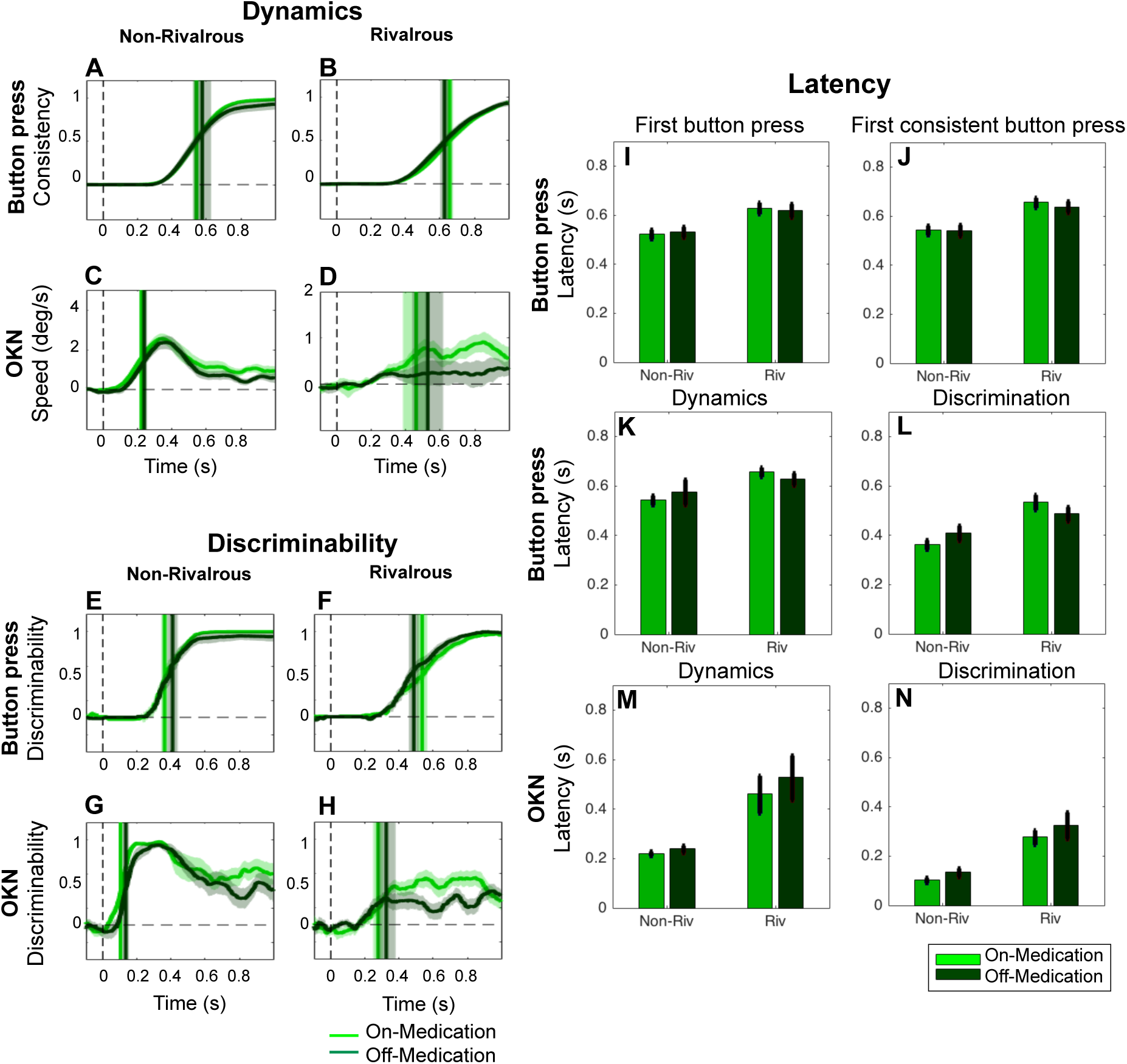
No effects of medication on button press and OKN (within subject comparison). The format of panel A-D, E-H, and I-N is the same as for Fig 2, 4, and 5. N=15 subjects were tested in the on-(light green) and off-(dark green) medication state. Note that y-axis for C and D are on different scales. No significant differences were found (FDR q=0.05 for corrections of multiple comparisons across time for A-H).

Table 2 summarizes subject demographics for each group in the first set of analysis. Although rejection rate of PD patients was significantly higher than controls (chi^2^=5.28, p=0.022), any particular rejection criteria did not differ significantly between the groups (button press: chi^2^=1.17, p=0.28; eye movements: chi^2^=0.65, p=0.42; both: chi^2^=3.56, p=0.6; imbalance: chi^2^=2.43, p=0.12).

**Table 2.** Demographics for subjects and rejected subjects for the first set of analysis, which compared controls and PD patients in on-treatment session. For the explanation of row 2 (rejected subjects), see Table 1. One PD subject is excluded due to missing MDS-UPDRS score in on-DBS session.

| Subject Group | Control | PD On-treatment |
| --- | --- | --- |
| Analyzed subjects | 24 | 22 |
| Age | 65.0(7.7 SD) | 66.5(8.0 SD) |
| MDS-UPDRS | NA | 13.6(7.6%) |

### PD patients can report their percepts with buttons as accurately as, but more slowly than controls

First, we examined the time of button press reports of the direction of perceived direction of motion in non-rivalrous conditions. Patients reported the correct direction of the physical stimulus as accurately as PD and controls (patients: M =92.5%, SEM=0.93; controls: M=93.6 %, SEM=0.55;unpaired two-tailed t-test: t(44)=0.98, p=0.32). However reaction times, measured as the first button press times, were prolonged in patients compared to controls (patients M=536 ms, SEM=16.7; controls: M=482 ms, SEM=13.3, unpaired two-tailed t-test: t(44) = −2.55, p=0.014).

Next, we examined the time of button press reports in rivalrous condition. While we cannot define the accuracy for rivalrous condition as we did for non-rivalrous condition, we can compare two types of button latency in patients and controls: the latency to first button press and the latency to first ‘consistent’ button press (as defined in Methods). The mean latency of first button press in rivalrous condition did not differ between the groups (patients: M=636 ms, SEM=24.8; controls: M=579 ms, SEM=18.8; unpaired two-tailed t-test: t(44)=-1.85, p=0.071). However, button press reports can switch during a 2 s rivalrous trial. Such a switch may reflect a genuine change, or a mistake — an initial impulsive (and brief) button press followed by a second corrective (and more prolonged) button press that reflects their genuine percept in that trial. Taking into consideration of these cases, we analyzed the latency of first ‘consistent’ button press (see Method). The mean latency of first consistent button press in rivalrous condition was significantly slower for PD patients than for controls (patients: M=655 ms, SEM=21.6; controls: M=596 ms, SEM=18.0; unpaired two-tailed t-test: t(44)=-2.13, p=0.039).

Taken together, we conclude that PD patients can respond as accurately as, but more slowly, than controls with button press in non-rivalrous condition, where we can objectively define the correct answer. In rivalrous condition, we cannot quantify the accuracy of button press, but PD patients started showing slower consistent responses than controls.

### Temporal dynamics of button press reports and OKN

Next, we analyzed the mean time course of button press reports as well as the velocity of the slow-phase OKN (Fig 2).

For button press reports, we analyzed whether button press was consistent with the labelled direction of the trial over time, in non-rivalrous and rivalrous conditions. Using this method, we confirmed that button press in PD is significantly slower than in controls in both non-rivalrous (Fig 2A) and rivalrous (Fig 2B) conditions.

Although the mean button press consistency in patients appears lower than that of controls between 0.4-0.8 s for both rivalrous and non-rivalrous condition (Fig 2A and B), the differences did not reach significance (unpaired two-tailed t-test at each time point, multiple comparisons were corrected with False Discovery Rate at q=0.05). Due to the smoothing procedure, we expected that the first consistent button press time would roughly correspond to when consistency reached 0.5, and this was confirmed to be the case. PD was significantly slower in reaching 0.5 in button press consistency than control for both non-rivalrous (unpaired two-tailed t-test: t(44)=-2.24, p=0.030) and rivalrous condition (unpaired two-tailed t-test: t(44)=-2.23, p=0.030).

Fig 2C and D show the mean time course of the velocity of slow-phase OKN. Note that we flipped the sign of the OKN for the left trials which tend to have negative velocity and that we scaled the y-axis differently for Fig 2C and D. Patients had a slower OKN than controls in both non-rivalrous (about 1.5 dg/s slower in PD, p<0.05 from 0.15 s to 2.0 s) and rivalrous conditions (about 1 dg/s slower in PD, p<0.05 at some points, as indicated by black lines at the bottom of Fig 2D, in 0.15 s to 2.0 s; unpaired two-tailed t-test at each time point, with FDR q=0.05).

Analogous to the latency for button press consistency to reach 0.5 (Fig 2A and B, vertical lines), we defined the latency of OKN as the time taken for OKN speed to reach the half of the maximum. The OKN latencies defined this way (Fig 2C and D vertical lines) were much faster than the button press latencies, which was around 0.5-0.6 s, in both patients and controls and in both stimulus conditions. We confirmed this with 3-way ANOVA of latency (defined as the time to reach half-maxima); we found significant main effects for the measure (button press vs OKN; F=97.00, p<0.001), the stimulus condition (rivalrous vs non-rivalrous; F=40.80, p<0.001) and the subject group (PD patients vs controls; F=6.33, p<0.05). As can be seen from Fig 2, OKN latency is much faster than button press especially in non-rivalrous condition, which is confirmed by a significant interaction between the measure (button press vs OKN) and the stimulus condition (rivalrous vs non-rivalrous) (F=7.6, p<0.01). Other interactions were not significant (stimulus*subject group: F=0.32, p=0.57; subject group*data type: F=0.27, p=0.60; stimulus*subject group*data type: F=0.75, p=0.39).

While OKN has a relatively short latency, its reliability in terms of discriminating the direction of perceived motion cannot be inferred from the mean time course. Thus, we turn our focus to trial-by-trial analysis of OKN to quantify how accurately OKN reflects conscious perception over time in each trial.

### Trial-by-trial analysis of button press and OKN

To quantify how accurately the initial OKN reflects the (non-rivalrous) stimuli or (rivalrous) percepts, we performed image-based trial-by-trial analysis of OKN and button press. Within each subject, we first sorted the trials by a mean button-press consistency value (over each 2-s trial) and in descending order, combined them to to form a mean button press consistency x time image for each subject. We then stretched each subject’s image to a uniform height, upsampled and then, smoothed. Note that we performed stretching, upsampling, and smoothing only along y-dimension, but not across time. Finally we averaged the images across subjects to obtain the mean button press consistency x time image for each group. (see Fig 3A).

In the mean button-press consistency x time images (Fig 3B, C, F and G), the color of each pixel represents the button press consistency at a particular time averaged across subjects. The upper rows in each panel represent trials with higher button-press consistency, which tend to have a shorter latency to first consistent button press; the lower rows represent trials with lower button-press consistency, which tend to have a longer latency to first consistent button press and have less time remaining to hold down that button before the trial ends. In this format, perceptual switches in rivalrous condition (Fig 4F and G) are captured by the lower consistency value towards the end of the trials.

Two important insights about OKN emerge from the image-based trial-by-trial analysis (Note the sign of OKN for the left trials are flipped as in Fig 2). First, OKN latency appears to be shorter and more uniform across trials compared to button press in both subject groups and stimulus conditions. This corroborates the observation in Fig 2. Second, OKN speed (depicted in color in each pixel) is quite variable across trials. To quantitatively compare the latency and variability of OKN with respect to those of button press consistency, we turn to the ideal observer analysis, next.

### Ideal observer analysis: discriminability of direction of stimulus and perceived motion based on the momentary button press and OKN

Taking into consideration OKN’s variable speed but shorter and more uniform latency compared to button press, we employed ideal observer analysis to quantify how reliably the velocity of slow phase OKN discriminates the direction of the stimulus (non-rivalrous condition) or conscious perception (rivalrous condition) in patients and controls (see Method).

We first computed the ability of button press to discriminate direction of non-rivalrous stimuli using the button press consistency (Fig 4A), which is a continuous measure after smoothing button presses over 0.1 s — the same amount of smoothing applied to OKN. The maximum of mean discriminability based on the button press consistency reached ~98 % by 0.5 s (where chance performance is 50 % and perfect performance is 100 %; The maximum discrimination did not reach 100 % for PD because one subject did not press the button correctly in some trials). Patients took longer than controls to reach half maximum discriminability (=~75 %) (vertical lines in Fig 4A) (unpaired two-tailed t-tests: t(44)=-3.49, p=0.001).

In rivalrous condition (Fig 4B), we labelled trials according to the dominant direction of perceived motion according to button press report during the first 1.5 s (see Methods). We computed the ability to discriminate the labelled direction of the trial based on button press consistency value at any one moment. The mean discriminability reached 100 % by 1 s. In contrast to non-rivalrous condition, the time to reach the half-maxima did not differ between patients and controls (unpaired two-tailed t-tests: t(44)=-1.15, p=0.258).

Applying the same analysis to OKN in non-rivalrous condition (Fig 4C), we found the maximum of mean discriminability of the OKN measure reached above 96 % within 0.2 s for both patients and controls. Interestingly, and in accord with previous analyses (Fig 2C, 3D and E), the speed of OKN decreased and became more variable across trials after 0.4 s, especially in the PD group. OKN-based discriminability degraded to ~70% in PD and while remained at ~90% in controls in the 0.5-0.8 s interval (p<0.05 with FDR q =0.05, black lines at the bottom of Fig 4C).

In rivalrous condition (Fig 4D), OKN-based discriminability was reduced (patients: ~70 %; controls: around ~80-90 %) compared with non-rivalrous condition when we average across subjects. The maximum discriminability was also lower for patients than controls’ (patients: M=89.1 %, SEM=2.1; controls: M=95.4 %, SEM=1.7; unpaired two-tailed t-test: t(44)=2.35 p=0.023). Comparing relative discriminability of OKN to button press consistency, we found that in non-rivalrous condition (Fig 4A and C), OKN-based discrimination reached to half of the maximum ~0.15 s faster than button press in both patients (paired two-tailed t-tests: t(21)=12.83, p<0.0001) and controls (paired two-tailed t-tests: t(23)=14.10, p<0.0001). Similarly, in rivalrous condition (Fig 4B and D), we found that OKN discriminability reached to half of the maximum faster than button press discriminability in both patients (paired two-tailed t-tests: t(21)=3.57, p=0.0018) and controls (paired two-tailed t-tests: t(23)=5.85, p<0.0001).

Taken together, in non-rivalrous condition, we conclude that OKN reflects the direction of the physical stimuli much faster than button press at a comparable accuracy. In rivalrous condition as well, OKN discrimination reached half of the maximum achievable faster than button press.

## Summary of Results

Fig 5 summarizes the results of button press and OKN for each subject group and stimulus condition. OKN (Fig 5E and F) has a shorter latency than button press (Fig 5A-D) across all all analysis methods for both subject groups and stimulus conditions.

We expected PD would delay motor responses across all motor response modalities, from voluntary button press to involuntary oculomotor reflexes. However, it is notable that on OKN-based measures, patients were significantly slower than controls on the OKN-based discriminability measure only in non-rivalrous condition.

Next we examine the effect of anti-Parkinson medication and deep brain stimulation on OKN and button press.

### No effects of anti-Parkinson medication on button press and OKN in both non-rivalrous and rivalrous condition

Dopaminergic medication, such as L-DOPA, reduces motor symptoms of PD, including tremor and rigidity. To investigate effects of medication on button press and OKN in non-rivalrous and rivalrous conditions, we examined 15 medically-treated (i.e. non-surgically treated) PD patients (a subset of PD patients included for the analyses so far) in defined on- and off-medication states. On-medication was defined as taking their usual anti-Parkinson medication, while off-medication was defined as after withdrawal of all anti-Parkinson drugs for at least 12 hours. Of the 22 patients tested on and off medication, 7 were rejected based on the rejection criteria detailed in Methods. Table 3 lists patient demographic and the proportion of subjects rejected due to criteria related to button press and eye tracking. As expected MDS-UPDRS motor disability scores were significantly improved by medication (on-medication: M=13.9, SD=8.4; off-medication: M=21.8, SD=11.4; paired two-tailed t-test: t(14)=-3.59, p<0.01).

**Table 3.**
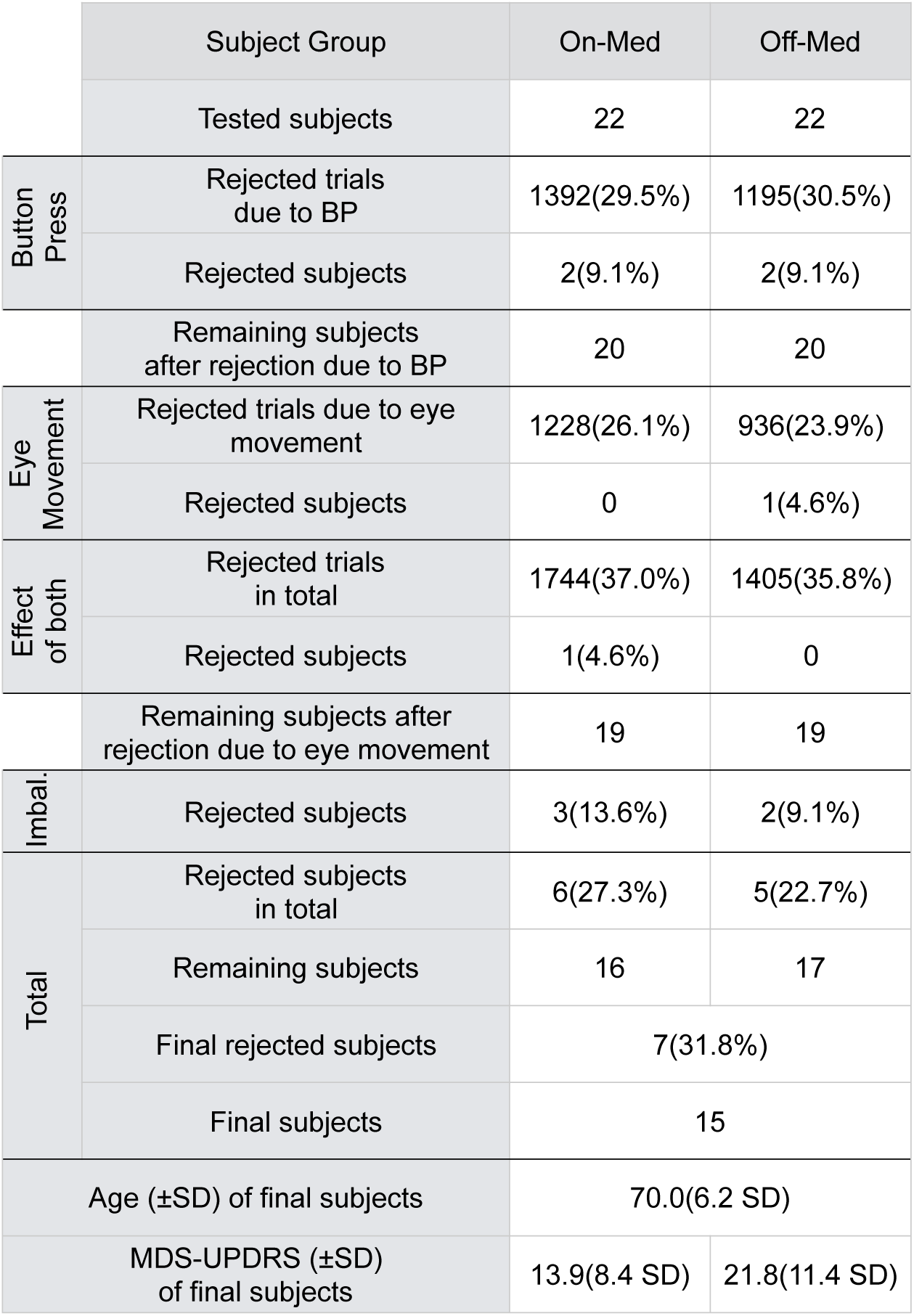
Subject demographics and rejected trials and subjects in the analyses comparing PD patients on and off medication. Trials were rejected due to the criteria for button press and eye movements (Table 1) that were explained in Methods. Subjects were rejected if more than half their trials in either of their on or off sessions was rejected. In total, we rejected 7 subjects (4 patients were rejected both sessions while the other 3 patients were rejected in one of the sessions).

|  | Subject Group | On-Med | Off-Med |
| --- | --- | --- | --- |
|  | Tested subjects | 22 | 22 |
| Button Press | Rejected trials due to BP | 1392(29.5%) | 1195(30.5%) |
|  | Rejected subjects | 2(9.1%) | 2(9.1%) |
|  | Remaining subjects after rejection due to BP | 20 | 20 |
| Eye Movement | Rejected trials due to eye movement | 1228(26.1%) | 936(23.9%) |
|  | Rejected subjects | 0 | 1(4.6%) |
| Effect of both | Rejected trials in total | 1744(37.0%) | 1405(35.8%) |
|  | Rejected subjects | 1(4.6%) | 0 |
|  | Remaining subjects after rejection due to eye movement | 19 | 19 |
| Imbal. | Rejected subjects | 3(13.6%) | 2(9.1%) |
| Total | Rejected subjects in total | 6(27.3%) | 5(22.7%) |
|  | Remaining subjects | 16 | 17 |
|  | Final rejected subjects | 7(31.8%) |  |
|  | Final subjects | 15 |  |
| Age (±SD) of final subjects | | 70.0(6.2 SD) | |
| MDS-UPDRS (±SD) of final subjects | | 13.9(8.4 SD) | 21.8(11.4 SD) |

For these analyses, we used the same procedure described in the patient vs control comparison (Fig 5) to compare patients on and off medication, except that we used paired t-tests in place of un-paired t-tests. As is clear from Fig 6, we found no effect of medication on either button press or OKN, despite the improvement in MDS-UPDRS motor disability scores.

### DBS facilitates button press in rivalrous condition, but did not affect on OKN and button press in non-rivalrous condition

Severe Parkinsonian symptoms, inadequately controlled by medication, can be effectively treated with Deep Brain Stimulation (DBS) (26). To investigate effects of DBS on button press and OKN in non-rivalrous and rivalrous conditions, we tested 17 patients (a subset of PD patients included in the initial analysis) on and off-DBS states. In the on-DBS condition, DBS settings were the patients’ usual therapeutic settings, while in the DBS-off condition, stimulation was turned off for at least 30 minutes before commencement of testing. Twelve of the DBS patients were also taking anti-Parkinson medication but there was no manipulation of their anti-Parkinson drugs.

Of the 17 PD patients tested on and off-DBS, 10 were rejected, leaving only 7 subjects in the final analysis. Table 4 lists the proportion of trials and subjects rejected due to to button press and eye tracking criteria. More subjects were rejected in off-state; 3 patients met rejection criteria in both on-DBS and off-DBS sessions, an additional 6 patients met rejection criteria only in the off-DBS session, and one other patient met rejection criteria in only the on-DBS session.

**Table 4.** Subject demographics and rejected trials and subjects in the analyses comparing PD patients on and off DBS. Trials were rejected due to the criteria for button press and eye movements (Table 1) that were explained in Methods. Subjects were rejected if more than half of trials in either of their on or off sessions was rejected. In total we rejected 10 subjects (3 were rejected both sessions, while the other 7 were rejected in one session).

|  | Subject Group | On-DBS | Off-DBS |
| --- | --- | --- | --- |
|  | Tested subjects | 17 | 17 |
| Button Press | Rejected trials due to BP | 333(26.9%) | 311(28.8%) |
|  | Rejected subjects | 2(11.8%) | 6(35.3%) |
|  | Remaining subjects after rejection due to BP | 15 | 11 |
| Eye Movement | Rejected trials due to eye movement | 347(28.0%) | 268(24.8%) |
|  | Rejected subjects | 1(5.9%) | 1(5.9%) |
| Effect of both | Rejected trials in total | 434(35.0%) | 386(35.7%) |
|  | Rejected subjects | 0 | 0 |
|  | Remaining subjects after rejection due to eye movement | 14 | 10 |
| Imbal. | Rejected subjects | 1(5.9%) | 2(11.8%) |
| Total | Rejected subjects in total | 4(23.5%) | 9(52.9%) |
|  | Remaining subjects | 13 | 8 |
|  | Final rejected subjects | 10(58.8%) |  |
|  | Final subjects | 7 |  |
| Age (±SD) of final subjects | | 66.5(8.0 SD) | |
| MDS-UPDRS (±SD) of final subjects | | 12.8(5.9 SD) | 30.2(9.7 SD) |

Fig 7 summarizes the results. We found that DBS reduced the latency of three button press related measures in rivalrous condition; latency to first button press (on-DBS: M=609 ms, SEM=51; off-DBS: M=675 ms, SEM=51; paired two-tailed t-test: t(6)=-4.20, p<0.01), latency to first consistent button press (on-DBS: M=637 ms, SEM=43; off-DBS: M=697 ms, SEM=44; paired two-tailed t-test: t(6)=-5.23, p<0.01) and the latency for button press consistency reach 0.75; on-DBS: M=642 ms, SEM=41; off-DBS: M=702, SEM=44 ms; paired two-tailed t-test: t(44)=-8.29, p<0.01).

**Fig 7.**
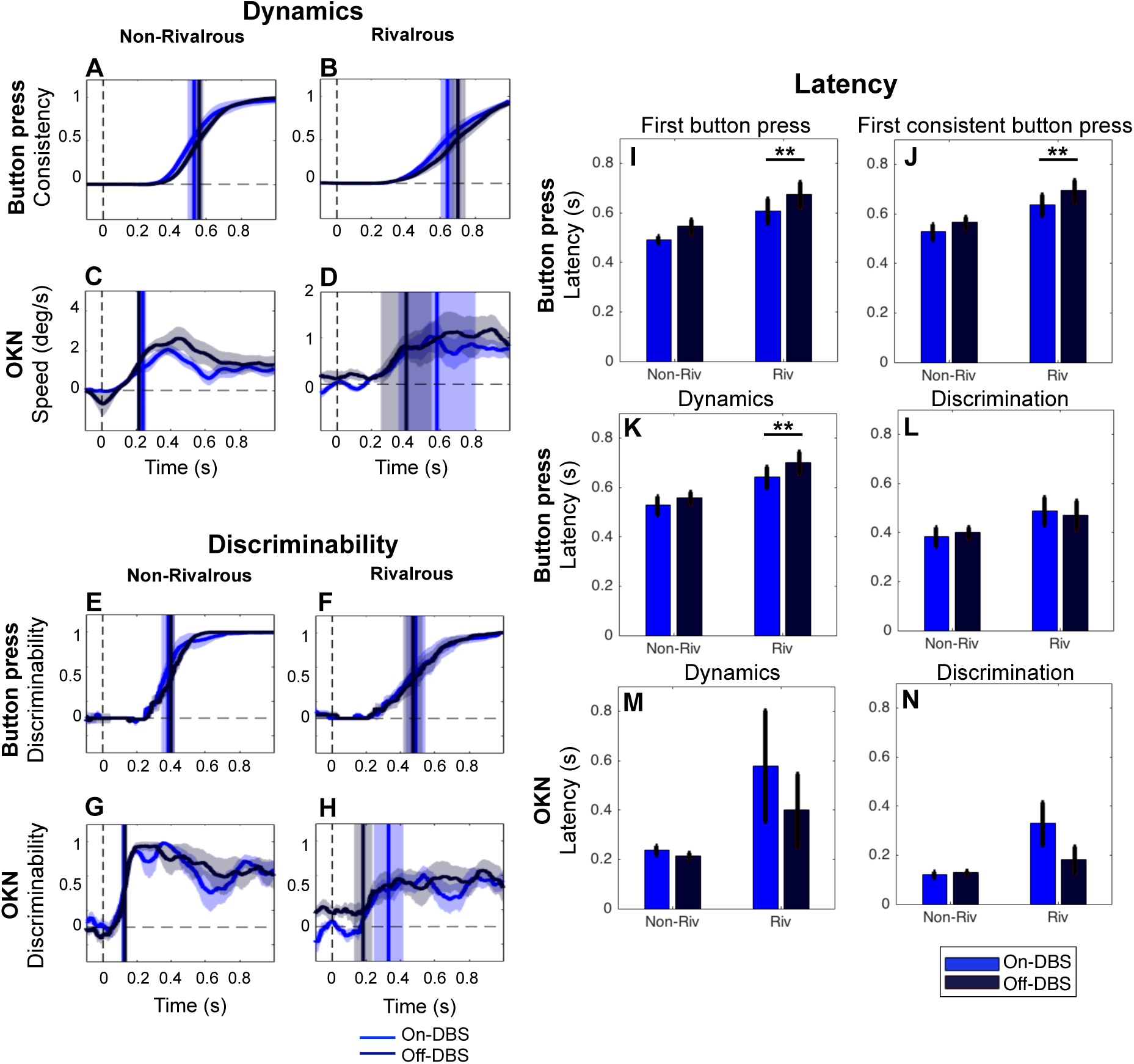
Deep brain stimulation facilitated button press but did not affect OKN. The format of the figure is the same as Fig 6. ** indicates significant difference (p<0.01). For A-H, corrections of multiple comparisons across time were performed with FDR at q=0.05.

### No difference between PD treated with medication or DBS

Finally, we examined if patients treated with DBS and medication differed in any aspects of the button press or OKN because baseline severity of disease may have been greater in the DBS group. Comparing MDS-UPDRS motor score between medication patients and DBS patients, both group of subjects were equally disabled in on-state and off-state (unpaired two-tailed t-test; on-state, t(19)=0.27, p=0.79; off-state, t(19)=-1.58, p=0.13).

Fig 8 compares PD patients treated with only medication (N=15, reported in Fig 6), and with DBS (N=7, reported in Fig 7). We averaged the data across on and off states in each subject, then performed the between-subject analysis with unpaired t-tests. The results did not differ between the groups in any of our button press or OKN measures.

**Fig 8.**
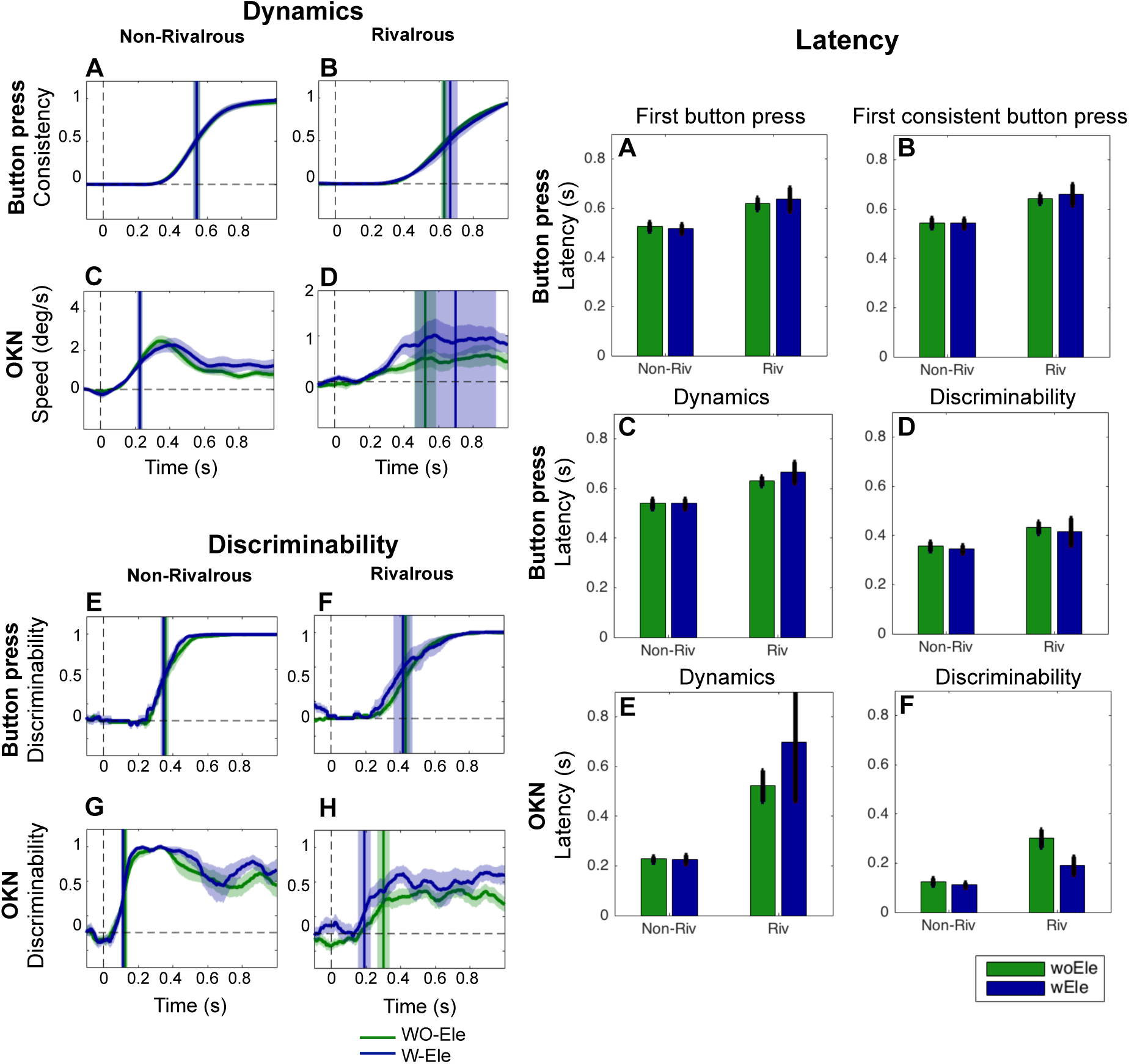
Treatment types of PD did not affect on our measures of button press and OKN. The format of the figure is the same as Fig 6 and 7. The N=15 medication patients and N=7 DBS patients were compared after averaging the data for two sessions for each subject. No significant differences were found.

## Discussion

We investigated conscious perception in PD patients using OKN as an additional measurement to behavioral responses. We found that OKN responses reflected perceptions in PD as accurately as in normal controls. OKN proved to be not only a reliable measurement of conscious perception but also faster at detecting perceived direction of motion than manual responses in both patients and controls. The speed of OKN was decreased in patients, but the latency of OKN temporal dynamics were not prolonged in comparison to controls. In contrast, button press latencies were prolonged in PD patients compared to controls, as expected due to their reduced mobility in PD.

Our findings have implications for psychophysical studies of PD patients. Psychophysical experiments performed on PD patients (e.g., (27)) have often employed verbal report by patients followed by delegated button press report by the experimenter to overcome presumed motor difficulties and other difficulties such as drowsiness (28). However, delegated response procedures are indirect and preclude measurement of reaction times. Consistent with previous psychophysical studies (29,30), our patients were also slower than controls in 6 out of 8 measures of button press in Fig 5A-D. However, the delay was not so dramatic to the extent that it invalidated the button press reports by the patients altogether. The degree of delay in button press seems to vary across patients and may depend on the difficulty of the psychophysical tasks.

### DBS facilitates button press response latency in rivalrous condition

Effect of DBS treatment was significant in 3 out of 4 measures related to button press in rivalrous condition (Fig 7). This result is consist with a proposal by Frank and colleagues (31) in which they suggested that DBS interferes with patients in their ability to slow down decision making under high-conflict conditions. It is plausible that perceptual conflict is induced in our rivalrous condition and that DBS specifically facilitates quicker decision in rivalrous condition. Interestingly, Frank and colleagues also reported that no such effect was found with dopaminergic medication. This too, is consistent with our observation of the absence of any effect of medication (Fig 6). While the general patterns of results are in agreement, the nature of conflicts seem quite different between our perceptual rivalry task and Frank’s cognitive conflict task. Further investigations are necessary to understand if DBS acts on the same neural mechanisms that are responsible for resolving perceptual and cognitive conflicts.

### Effects of PD on OKN

Previous reports on the effect of PD on the OKN are mixed; some studies found that the OKN is impaired in PD patients (6,7,9), while others report that their OKN is intact (2,3,21). In our study, patients had a slower OKN compared with controls, especially in non-rivalrous condition (Fig 2C). The lower velocity compromised calculation of our mean OKN-based discriminability measures; however, each individual’s OKN predicted the direction of stimulus (non-rivalrous condition) or perceptual motion (rivalrous condition) to a comparable degree between PD and controls, especially at the trial onset. After 0.4 s, however, the discriminability started to diverge between PD and control. It is plausible that perceptual rivalry as well as OKN may be supported by distinct neural mechanisms at the onset phase and subsequent continuous phase (22).We are currently investigating the effects of PD in a continuous rivalry paradigm.

Neither medication (Fig 6) nor DBS (Fig 7) affected much on the velocity, as well as the discriminability, of OKN. This suggests that impairment of OKN speed is a stable deficit and not modifiable by manipulating anti-Parkinson drugs or DBS, at least in the short term.

### OKN as a readout of conscious perception

OKN during binocular rivalry with moving stimuli is shown to correlate well with direction of consciously perceived perception in healthy populations (13–18,32). Here, we extended the generalizability of OKN readout in rivalry to patients with PD.

The intricacies of the relationship between OKN and conscious perception has not been fully elucidated. In fact, direction of eye movements is not always correlated with conscious perception (33). When orthogonal horizontal and vertical moving gratings are presented to both eyes separately, the eyes move in the average direction of the two gratings, while subjects only perceive either horizontal or vertical direction of motion, demonstrating dissociation of eye movements and percept (34). Unlike such a study, we used two gratings moving to the opposite directions, which has been shown to induce strong concordance between eye movements and conscious percept (11,18). Fully elucidating the conditions that promotes concordance or dissociation between eye movements and conscious percept, combined with neuronal imaging, will help facilitate our understanding of the neural mechanisms of consciousness. When OKN are concordant with conscious percept, as in our case, OKN can be used as a reliable readout of conscious perception without requiring button press reports (11,12,14,35). When OKN dissociates from reported conscious perception, OKN can be used to examine the behavioral and neural mechanisms of the non-conscious visual processing (33,34).

OKN has received strong attention recently as it can be used as a potential readout of conscious percept in no-report paradigms (12). No-report paradigms, combined with traditional report-based paradigms, allow researchers to separate neuronal activity generated by manually reporting from the neuronal activity of the percept itself. For example, Frassle and colleagues demonstrated that neural activity in the prefrontal cortex during binocular rivalry diminished strikingly under the no-report condition compared with voluntary report conditions. This and other convergent evidence from no-report paradigms (12,36) suggests that the activity in the prefrontal cortex may be more related to the act of reporting, and that the prefrontal cortex may not be a critical neuronal correlate of consciousness as suggested by studies that only included voluntary perceptual reports (37–40).

Our trial-by-trial analyses (Fig 3) in non-rivalrous condition poses an interesting question. There, in addition to being faster than button press, OKN showed less variability in response latency across trials. With our current methods, we cannot tell which measure (button press or OKN) correlates best with the latency of conscious perception of visual motion. Button press is susceptible to various factors, such as motor preparation and attention, with unknown effects on the latency of conscious perception (41). It is tempting to suggest that OKN’s faster and less variable response might better reflect the onset of conscious perception than the button response modality, which is currently the dominant measure in psychological studies of consciousness. Broadening the response modalities from momentary button press to continuous measures, such as OKN, may be more effective in capturing complex perceptual dynamics during rivalry (42,43) and in elucidating the gradual nature of consciousness and its neural basis (11).

### Study Limitations: Rejections of PD patients

A relatively high rejection rate of severely affected patients prompts cautious interpretation of some of our results.

Around 30 % of trials were rejected for both control and PD (Table 1), mainly because we strictly required subjects to release the button every 2 s after the stimulus period. This required considerable effort from subjects, but it was necessary to properly estimate the latency of button press.

A more interesting pattern of rejection is to do with perceptual imbalance (Table 1), because of which PD patients were more likely to be rejected than than controls, no controls and 6 patients were rejected, although the rejection rates were not statistically different (chi-square test, p=0.12). This implies that PD patients may be more likely to stick with one percept across many rivalrous trials. Reduced perceptual switches (also known as “perceptual freezing” or “perceptual memory”) under the intermittent presentation of ambiguous stimuli has been extensively studied in healthy subjects (44,45). It would be interesting to test whether PD patients show exaggerated perceptual stabilization. Such a study may provide clues to the critical neuronal loci responsible for perceptual stabilization, as well as perceptual switches — Einhäuser et al (46) have proposed the locus coeruleus as a potential neural locus for perceptual switches during ambiguous stimulation and that norepinephrine, a precursor of dopamine, may be involved. The locus coeruleus is affected early in PD (47). Future studies employing OKN readout with no report to study in perceptual stabilization in PD patients may be fruitful in elucidating perceptual consequence of depletion of dopamine in PD patients.

## Acknowledgements

We are very grateful to all subjects participated in this study. LK and NT were supported by the Australian Research Council (ARC) Discovery Project (DP130100194). LK was supported by a fellowship from the Japanese Society for the Promotion of Science (JSPS P15048). NT was supported by Precursory Research for Embryonic Science and Technology project from the JST (3630) and ARC Future Fellowship (FT120100619). CD is supported by the Monash Institute of Neurological Sciences and an Australian Postgraduate Award from Monash University. We also appreciate Megan O’Neill and Eloise Perini for their contributions to the preliminary version of this project.

